# Body size affects immune cell proportions in birds and non-volant mammals, but not bats

**DOI:** 10.1101/2020.12.18.423538

**Authors:** Emily Cornelius Ruhs, Daniel J. Becker, Samantha J. Oakey, Ololade Ogunsina, M. Brock Fenton, Nancy B. Simmons, Lynn B. Martin, Cynthia J. Downs

**Author notes:** Shared first authorship (Emily Cornelius Ruhs, Daniel J. Becker).

## Abstract

Powered flight has evolved several times in vertebrates and constrains morphology and physiology in ways that likely have shaped how organisms cope with infections. Some of these constraints likely have impacts on aspects of immunology, such that larger fliers might prioritize risk reduction and safety. Addressing how the evolution of flight may have driven relationships between body size and immunity could be particularly informative for understanding the propensity of some taxa to harbor many virulent and sometimes zoonotic pathogens without showing clinical disease. Here, we used a scaling framework to quantify scaling relationships between body mass and the proportions of two types of white blood cells--lymphocytes, and granulocytes (neutr-/heterophils)--across 60 bat species, 414 bird species, and 256 non-volant mammal species. By using phylogenetically-informed statistical models on field-collected data from wild Neotropical bats, data gleaned from other wild bats available in the literature, and data from captive non-volant mammals and birds, we show that lymphocyte and neutrophil proportions do not vary systematically with body mass among bats. In contrast, larger birds and non-volant mammals have disproportionately higher granulocyte proportions than expected for their body size. Future comparative studies of wild bats, birds, and non-volant mammals of similar body mass should aim to further differentiate evolutionary effects and other aspects of life history on immune defense.

**Summary statement:** Powered flight might constrain morphology such that certain immunological features are prioritized. We show that bats largely have similar cell proportions across body mass compared to strong allometric scaling relationships in birds and non-flying mammals.

## Introduction

Powered flight has evolved at least three times in the evolutionary history of vertebrates and yet is one of the most energetically costly modes of transportation (Rayner, 1988). Birds and bats experience a 6–14 fold and >25 fold increase over resting metabolic rate, respectively, in metabolic expenditure during flight, whereas a similarly-sized mammal only experiences a 6–8 fold increase during sustained running (Schmidt-Nielsen, 1972; Thomas, 1975). Although there is some debate over whether bats or birds are more efficient fliers (Muijres, Johansson, Bowlin, Winter, and Hedenström, 2012; Swartz et al., 2007; Tian et al., 2006), there are clear functional and physiological constraints associated with this costly activity (Maurer et al., 2004; Muijres et al., 2012). One of the most evident constraints is body size. Exceptionally large and small body sizes have apparently been selected against in the evolution of flying vertebrates due to demands imposed by the physics of flight (Stanley, 1973); however, the constraining factors for bats and birds likely differ, as the largest bats are much smaller than the largest flying birds. The evolution of flight and body size constraints may have had numerous direct and indirect effects on evolution of the immune system in flying vertebrates. For example, evolution of a lightened skeleton (Feduccia and Feduccia, 1999; Dumont, 2010) may affect how immune cells are differentiated and distributed throughout the body. If larger fliers are not as efficient at circulating protective cells throughout their bodies, then they might require greater quantities of cells (Ruhs et al., 2020). It should be noted that the high energetic costs of flight have varying impacts on the immune system (Hasselquist et al. 2007; Voigt et al. 2020; Nebel et al. 2012). While birds and bats have much in common in terms of constraints that accommodate the ability to fly, the evolution of flight likely impacted the dynamics between body size, physiological traits, and the exposure risk to pathogens relative to non-flying birds and mammals.

Body size influences almost all life processes and structures of organisms (Brown, Gillooly et al. 2004; West et al. 2000). Many biological traits vary with body size in predictable ways; some vary proportionally across body size (i.e., isometric scaling), whereas others change disproportionately with size (i.e., hyper or hypometric scaling; Calder, 1996; Kleiber and Others, 1932; Knut Schmidt-Nielsen and Knut, 1984). Most efforts to describe relationships between size and traits take the form:

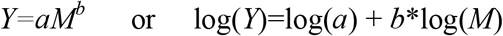

where (in the linearized form) *b* represents the scaling coefficient, *M* is body mass, *a* denotes the intercept, and *Y* represents the trait of interest. Many traits influenced by body size, including lifespan and movement patterns (e.g., home range size, distance traveled while foraging), affect pathogen exposure (Han et al., 2015), which could in turn exert selective pressure on how species allocate resources to immune defense (Brace et al., 2017; Lee, 2006).

Although various hypotheses predict distinct forms of scaling for aspects of immunity (Cohn and Langman, 1990; Dingli and Pacheco, 2006; Wiegel and Perelson, 2004), there is strong evidence that particular immune cells (namely concentrations of granulocytes, such as the neutrophils of mammals (Downs et al., 2020) and heterophils of birds (Ruhs et al. 2020) scale hypermetrically with body size. These allometries support the Safety Factor Hypothesis, which proposes that larger animals favor infection risk reduction by investing heavily in safety (Downs et al., 2020; Harrison, 2017), in particular by using a reserve pool of broadly protective granulocytes (e.g., neutrophils and heterophils). However, concentrations of heterophils in birds scale at a steeper rate (*b*=0.19; Ruhs et al., 2020) than mammals (*b*=0.11; Downs et al., 2020). Although this difference could be an evolutionary artifact or driven by one of many other differences between birds and mammals, this steep allometry has been hypothesized to be related to flight, which may put larger birds at a higher risk of parasite exposure (Downs et al., 2019; Ruhs et al., 2020). It should also be noted that larger birds and bats also have longer lifespans than similarly sized non-volant mammals (Munshi-South and Wilkinson, 2010; Wilkinson and Adams, 2019), possibly putting them at increased risk of infection over their long lifespans. The potential for larger fliers to prioritize risk-reduction immunological strategies motivates our interest to investigate immune scaling in bats and among bats, birds, and other mammals.

Bats are a hyperdiverse taxon (Order Chiroptera, over 1400 species) with a nearly global distribution across habitats ranging from rainforests to deserts (Gunnell and Simmons, 2012; Simmons and Cirranello, 2020). Their unique habits and life histories (e.g., powered flight, echolocation, long lifespans despite small body sizes) make bats a notable taxon for basic studies of ecology and evolution (Ingala et al., 2018; Jones and Teeling, 2006; Wilkinson and South, 2002). Bats have also been increasingly studied for their ability to harbour some viruses that are detrimental and often lethal to humans and domestic animals (Brook and Dobson, 2015; Guth et al., 2019). Bats are confirmed reservoir hosts for henipaviruses, Marburg virus, various lyssaviruses, and most SARS-like coronaviruses (Amman et al., 2015; Banyard et al., 2011; Halpin et al., 2011; Li et al., 2005). Yet with some exceptions (e.g., *Rabies lyssavirus*), these viruses appear to not kill and rarely cause clinical disease in bats (Williamson et al., 2000).

Whereas the high diversity of zoonotic viruses in Chiroptera might be partly driven by the speciose nature of this order (Mollentze and Streicker, 2020), bat tolerance of particular viruses may be shaped by specialized immune mechanisms in these flying mammals (Brook et al., 2020; Zhang et al., 2013). Bat immunoglobulins and leukocytes are structurally similar to those of humans and mice (Baker et al. 2013), but bats also have unique immune system traits such as complement proteins robust to temperature change, lack of fever with bacterial (lipopolysaccharide) challenge, high constitutive expression of type I interferons, and dampened inflammation (Ahn et al., 2019; Hatten et al., 1973; Pavlovich et al., 2018; Stockmaier et al., 2015; Zhou et al., 2016). Flight may explain these distinctions, including increased metabolic rates that enable stronger immune responses and elevated body temperature that could mirror febrile responses to control infection (O’Shea et al. 2014; but see Levesque et al. 2020). However, the primary hypothesis for how bats can tolerate viruses is that they evolved mechanisms to minimize or repair the negative effects of oxidative stress generated as a consequence of flight (Zhang et al., 2013). For example, some bat species show resistance to protein oxidation and unfolding (Salmon et al. 2009), reduced lipid peroxidation (Wilhelm Filho et al. 2007), and lower hydrogen peroxide per unit oxygen consumed (Brunet-Rossinni 2004). This propensity to resist acute oxidative stress and repair oxidative damage could have also helped bats cope with viral replication that would have otherwise caused cell damage (Kacprzyk et al., 2017; Xie et al., 2018; Zhang et al., 2013).

Here, we first asked whether leukocyte proportion scaling in bats is distinct compared to other taxa already described. Then, we asked whether the ability to fly (i.e. bats and birds) explains immune cell proportion allometries across extant vertebrate endotherms. Most studies assessing immunity in bats have been limited to few species (but see Schneeberger et al., 2013). We combined field-collected data from Neotropical bats with data from the primary literature to maximize sample sizes as well as phylogenetic and body size diversity. We then quantified scaling relationships for proportions of two primary leukocytes for which abundant data were available, lymphocytes and granulocytes. Lymphocytes include B and T cells, which provide specific, but time-lagged, protection through antibody production and coordination of cascading immune responses. Granulocytes (neutrophils in mammals and heterophils in birds) are phagocytes that rapidly protect against micro- and macroparasites without education or much specificity (Lanier, 2013). Finally, we directly compared scaling relationships for cell proportions in bats to those of birds and non-volant mammals using an existing database (ZIMS).

We predicted the forms of relationships between cell proportions and body size based on previously discovered scaling patterns among body size and cell concentrations (Downs et al., 2020; Ruhs et al., 2020). Proportions, however, sometimes pose difficulties for studies of allometry because they are bound rather than having no continuous upper limit and are inherently co-dependent (i.e. one cell type goes up, another goes down). As leukocyte concentration data for bats are extremely rare in the literature, and proportional data permitted comparisons that were otherwise presently impossible, we estimated scaling patterns using proportional data but encourage caution in comparing results from this analysis against prior scaling for leukocyte concentrations. We hypothesized isometry for lymphocytes in bats, as was observed previously for bird cell concentrations (Ruhs et al., 2020). We expected isometry to manifest because lymphocytes are a functionally diverse group of cells including both B and T cells (Lanier, 2013), the proportions of which could vary dramatically among species. By contrast, granulocyte functions are fairly homogeneous, so we predicted hypermetric granulocyte scaling in bats as was observed in other mammals and birds (Downs et al., 2020; Ruhs et al., 2020). However, as larger fliers might overinvest in safety, we expected bat neutrophil proportions to scale hypermetrically, to the same degree (steeper than non-volant mammals) as was observed for heterophil concentrations in birds (predictions based on if flight influences scaling; Fig. 1; Ruhs et al., 2020). Alternatively, we could observe no impacts of flight on cell proportion allometries, which could be due to life-history features (e.g. reproduction, sociality) or equal investment in risk reduction strategy due to factors like increased lifespan, regardless of body size.

**Figure 1.**
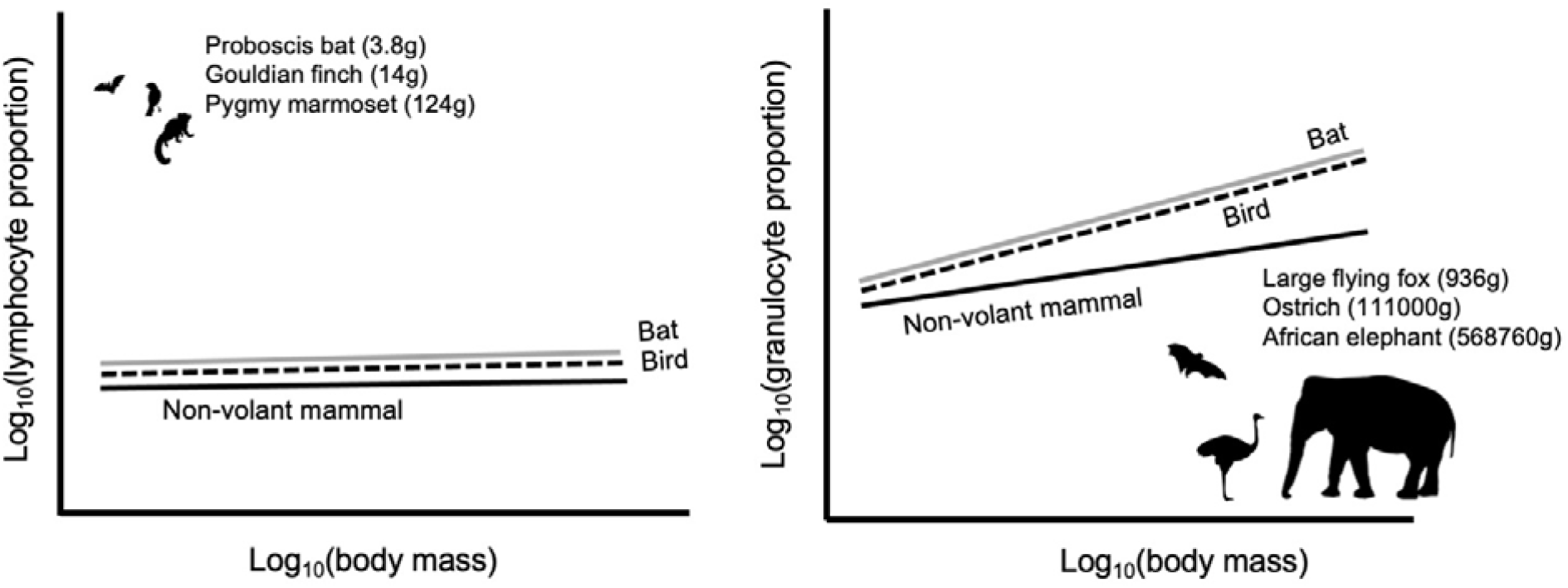
Predictions based on flight influencing the scaling relationship between (left) lymphocyte and (right) granulocyte proportions and body mass. Animals in the figures represent the smallest and largest animals in the datasets. For the rationale of our predictions, please see the main text.

## Materials and methods

### Bat sampling

During April and May in 2017 and 2018, we sampled 160 bats from 26 species in the Orange Walk District of Belize (Herrera et al., 2018). Bats were captured using mist nets and harp traps and were identified to species based on morphology (Reid, 1997). We collected blood by lancing the propatagial vein with a sterile needle, followed by collection using heparinized capillary tubes. Thin blood smears were prepared and stained with buffered Wright–Giemsa (Astral Diagnostics Quick III). All bats were released after processing. Sampling followed guidelines for safe and humane handling of bats from the American Society of Mammalogists (Sikes and Gannon, 2011) and was approved by the Institutional Animal Care and Use Committees of the University of Georgia (A2014 04□016□Y3□A5) and American Museum of Natural History (AMNHIACUC□20170403). Sampling was authorized by the Belize Forest Department under permits WL/2/1/17(16), WL/2/1/17(19), WL/2/1/18(16).

### Bat leukocyte data

We used light microscopy (1000X) to quantify the proportion of neutrophils and lymphocytes from 100 leukocytes from each field sample (Schneeberger et al., 2013). As Neotropical bats are relatively limited in their range of body masses, we supplemented our leukocyte dataset with a systematic literature search (Fig. S1). We identified articles using Web of Science and the search terms TS=(bat OR Chiroptera OR flying fox) AND (hematology OR white blood cell OR leukocyte). For bat species sampled across multiple studies, we averaged cell proportions. When available, body masses of each bat species were extracted from EltonTraits (Wilman et al., 2014); however, for a few species (*n*=2), masses were averaged from the source paper. The literature search substantially increased our body mass range (from approximately 5-78 grams to 4-804 grams; see Figure S2 for a comparison across all extant bat species; Wilman et al., 2014). Within-species sample size ranged from 1 to 140 (x□=15 ± 3) but did not predict proportions of either cell type (neutrophils: ρ=-0.09, *p* = 0.48; lymphocytes: ρ=0.06,*p*=0.67).

### Bird and non-volant mammal data

To compare bat leukocyte data to comparable data from birds and non-volant mammals, we extracted species means of lymphocyte and granulocytes (neutr-/heterophils) proportions in whole blood from ZIMS (Species360, 2019). ZIMS is a repository of veterinary data from captive, adult animals housed in facilities accredited by the Association of Zoos and Aquariums and considered healthy. We removed bat data (*n*=3) from the extracted ZIMS mammal database. When cleaning the data, we only included data from Global Species Reference Intervals. We compiled standardized species-level body mass data from the CRC Handbook of Avian Masses (Dunning Jr., 2007) and/or publicly available databases such as AnAge (Tacutu et al., 2013), the Animal Diversity Website (Jones et al., 1997), and the Encyclopedia of Life (Parr et al., 2014).

### Statistical analyses

#### Exercise 1: best-fit models for leukocyte proportion allometries in birds, bats, and non-volant mammals

Our modeling progressed in two stages. First, to test hypotheses about allometric scaling of leukocytes in bats only, we used phylogenetic generalized mixed-effects models (GLMMs) with the *ape* and *MCMCglmm* packages in R (Hadfield & Others, 2010; Paradis et al., 2004). All models included phylogenetic effects from a phylogeny produced in PhyloT using data from the National Center for Biotechnology Information (Letunic, 2015) and with resolved polytomies. We used that tree to create two phylogenetic covariance matrices, one for bat-only analyses and one that we used later for direct comparisons of scaling slopes across taxa. We set the inverse-gamma priors to 0.01 for the random effect of phylogenetic variance and default priors for the fixed effects in all models. All models were run for 260k iterations with 60k burn-in and a 200-iteration thinning interval (Downs et al., 2020; Ruhs et al., 2020). For all models, we used Deviance Information Criterion (ΔDIC) to identify the best-fit GLMM. We defined the top model as that with the lowest DIC, and we considered models within ΔDIC<5 as having equivalent support (Richards, 2005). For all models, we also calculated Pagel’s unadjusted λ and conditional and marginal *R^2^* (Housworth et al., 2004; Nakagawa and Schielzeth, 2013). We then used this approach to determine the scaling relationship for lymphocyte and neutrophil proportions, separately, across 60 bat species. Also, because previously published slopes for mammal and bird leukocyte scaling used cell concentrations (Downs et al., 2020; Ruhs et al., 2020), we determined the scaling relationships of cell proportion data for both birds (*n*=414) and mammals (*n*=256) independently to facilitate direct comparisons with bats. Results from bird and mammal-only models are presented in the online supplement (Tables S1 and S2). For each taxonomic group and each cell type, we produced two sets of *a priori* models:

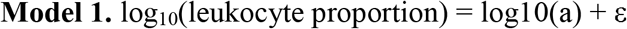

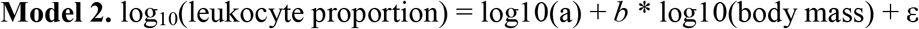

Model 1 represents a null model with *b*=0, whereas model 2 estimates the scaling relationship between body mass and cell proportion.

#### Exercise 2: direct comparisons of allometries among taxa

Next, we directly compared the slopes of relationships between body mass and immune cell type in bats (*n*=60), birds (*n*=414), and non-flying mammals (*n*=256). Specifically, we fit five models to the data and compared DIC scores to determine the best-fit versions

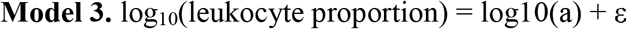

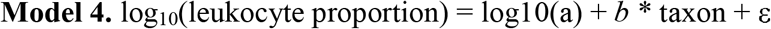

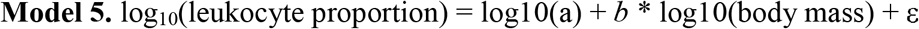

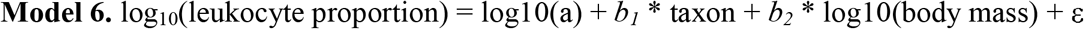

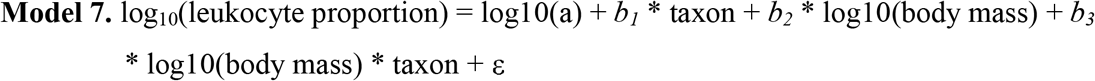

Here, model 3 represents a null model with *b=0,* and model 4 only tests for mean differences in cell proportions per taxon (i.e., bat, bird, non-volant mammal), irrespective of body mass. Model 5 is analogous to model 2 from exercise 1, and it estimates a global scaling relationship between body mass and cell proportions across all taxa. Lastly, model 6 combines models 4 and 5 (i.e., mean differences between taxa and a global body mass slope), whereas model 7 explicitly tests whether scaling relationships between body mass and cell proportions differ among taxa.

## Results

### Exercise 1: best-fit models for leukocyte proportion allometries in birds, bats, and non-volant mammals

For bats, the intercept-only model (fitting *b*=0; model 1) and the mass model (model 2) received equivalent support for both lymphocytes and neutrophils (Table 1, Fig. 2). Slopes for lymphocytes (*b*, CI=-0.08, −0.24:0.08) and neutrophils (*b*, CI=0.06, −0.09:0.22) were indistinguishable from zero (Table S1, Figs. 2 and 3). Although these results suggest isometry for lymphocytes and neutrophils, we encourage caution in their interpretation given that null models can represent either a true slope of zero or a lack of power to find allometry. For lymphocytes, phylogeny accounted for very little of the variation (16%) and the addition of body mass increased explanatory power by 15%. For neutrophils, again, phylogeny accounted for very little of the variation (21%), and the addition of body mass increased explanatory power by only 4%. For both leukocyte types, the model fit of the mass model was 33-37% and did not increase much from the 19-29% of the intercept-only model. Collectively, our results suggest little allometric scaling of proportions of either cell type among bat species.

**Table 1.** Best-fit models predicting circulating leukocyte concentrations in 60 species of bats (exercise 1). Models test for the effects of body mass on log_10_-transformed lymphocyte and neutrophil proportions. For all models, we calculated (1) Pagel’s unadjusted lambda to determine the variation explained by the phylogeny not accounting for fixed effects, (2) marginal R^2^ values to determine how much variation in leukocyte concentrations was explained by fixed effects and (3) conditional R^2^ for overall model fit.

| Model | DIC | ΔDIC | λ (unadjusted)<br>[95% CI] | Marginal<br>R<sup>2</sup><br>[95% CI] | Conditional<br>R<sup>2</sup><br>[95% CI] |
| --- | --- | --- | --- | --- | --- |
| <u>Lymphocytes</u> |  |  |  |  |  |
| 1. $\beta_0$ | 4.12 | 0.15 | 0.19<br>[0.04-0.62] | | 0.19<br>[0.04:0.62] |
| 2. $\beta_0 + \beta_1 \times \log_{10}(\text{Mass})$ | 3.96 | 0 | 0.16<br>[0.06:0.63] | 0.002<br>[2.1e-6:0.17] | 0.33<br>[0.07:0.68] |
| <u>Neutrophils</u> |  |  |  |  |  |
| 1. $\beta_0$ | -5.88 | 0 | 0.29<br>[0.07-0.68] | | 0.29<br>[0.07:0.68] |
| 2. $\beta_0 + \beta_1 \times \log_{10}(\text{Mass})$ | -5.28 | 0.6 | 0.21<br>[0.06:0.69] | 0.001<br>[1.2e-7:0.15] | 0.37<br>[0.09:0.72] |

**Figure 2.**
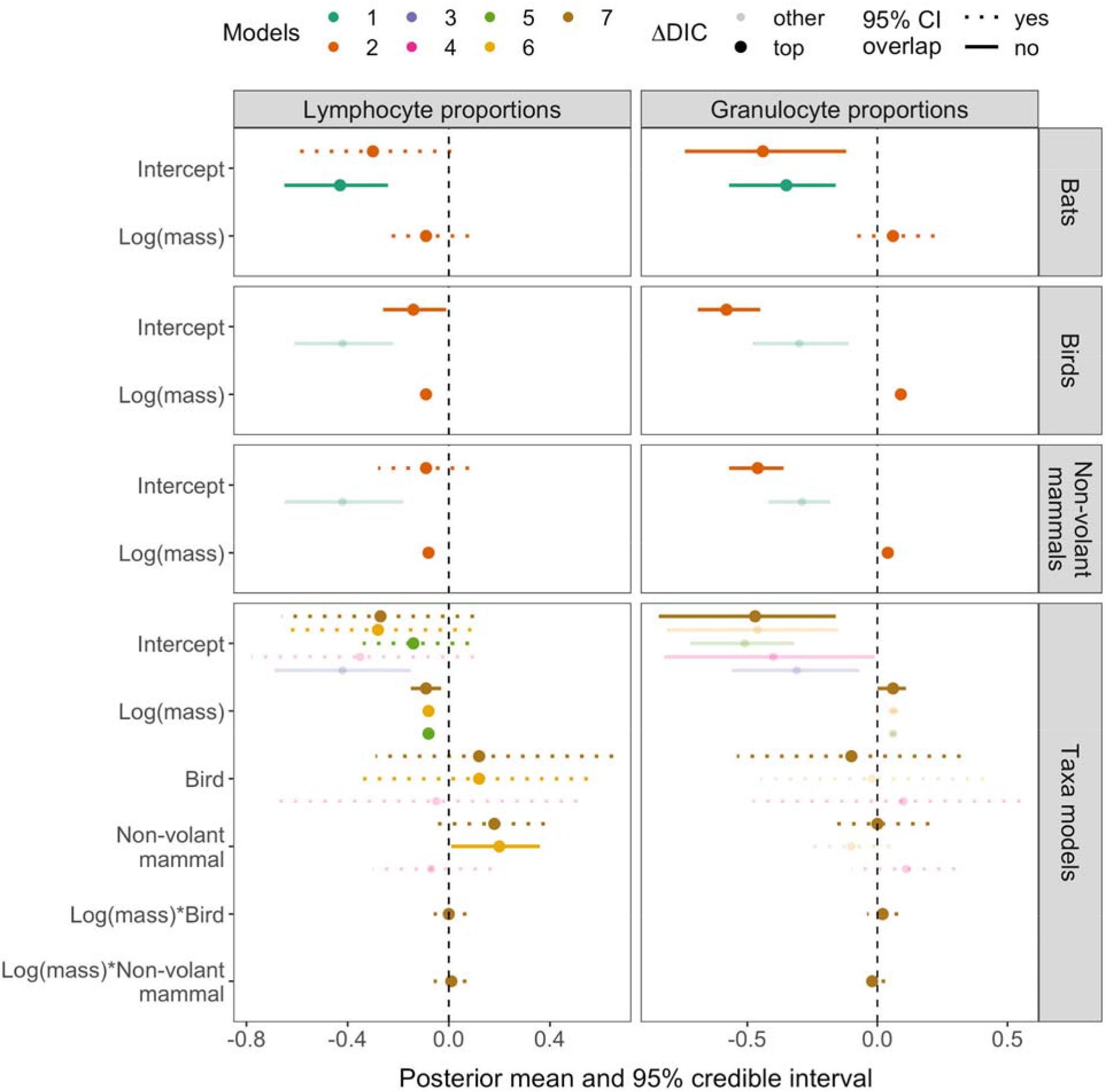
Posterior means and 95% credible intervals for coefficients of allometric scaling models applied to proportions of lymphocytes and granulocytes (neutrophils or heterophils) per each taxon and compared across taxa. Results are highlighted from the top models (ΔDIC<5, indicated by point size), whereas other competing models are transparent. Credible intervals that do not overlap with zero are displayed with solid lines. In the models comparing taxa, bats are represented by the intercept (models 4, 6, 7).

**Figure 3.**
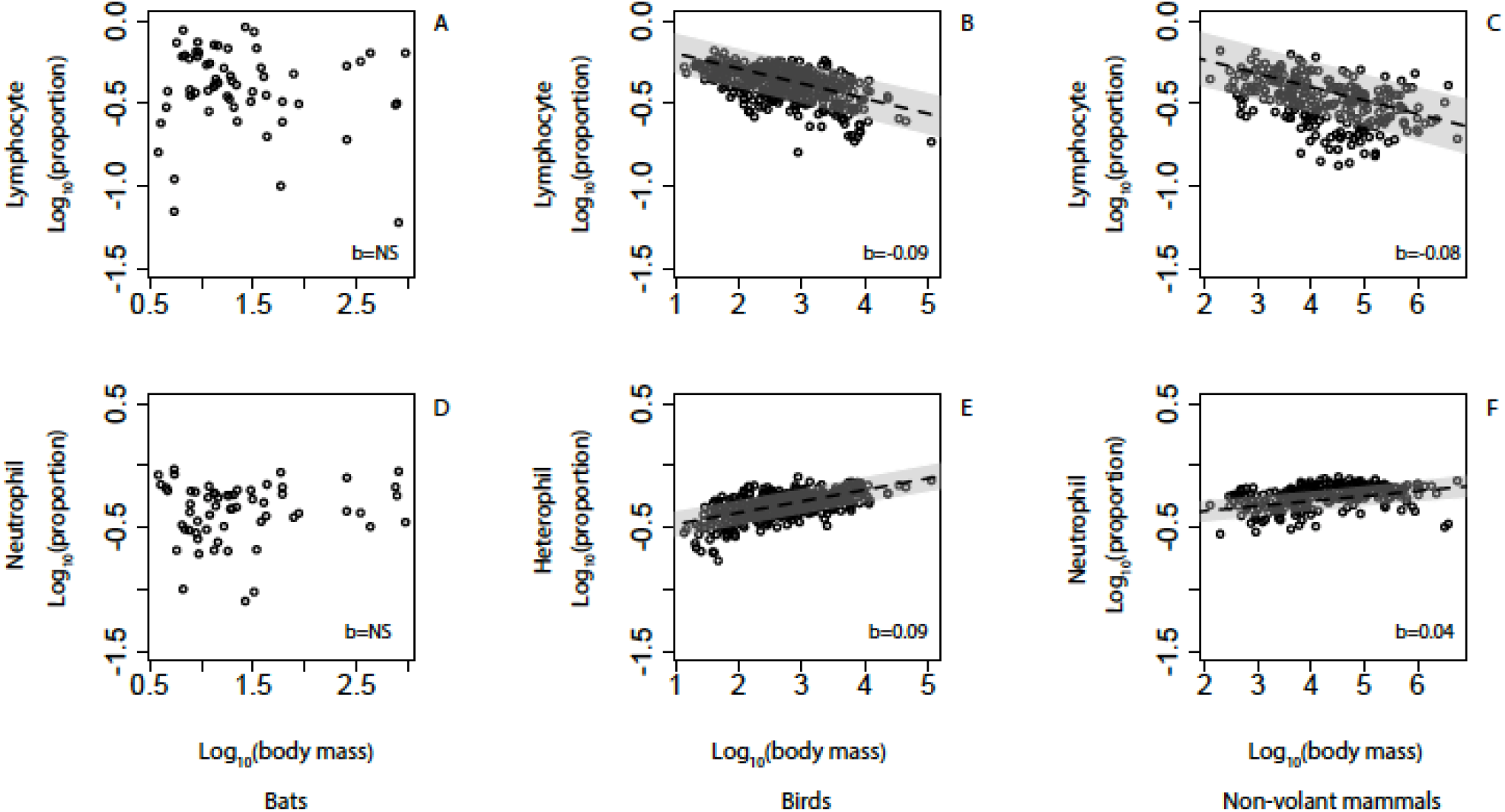
Observed scaling relationships in all species (bats (*n*=60), birds (*n*=414), and nonflying mammals (*n*=256)) between body mass and (A-C) lymphocyte and (D-F) neutr-/heterophil proportions. Dotted lines depict 95% credible intervals of the slope estimates. Data are plotted from model 2 (mass inclusive) from exercise 1.

For birds, the mass model (model 2) was best-supported (Table S2) for both cell types; lymphocytes scaled hypometrically (*b,* CI=-0.09, −0.1:-0.08; Table S3; Fig. 3) and heterophils scaled hypermetrically (*b,* CI=0.09, 0.08:0.11; Table S3). For non-volant mammals, the lymphocyte and neutrophil mass models were also the best-supported (Table S4); lymphocytes scaled hypometrically (*b*, CI=-0.08, −0.1:-0.06; Table S5; Fig. 3) and neutrophils scaled hypermetrically (*b*, CI=0.04, 0.03:0.06; Table S5). Phylogeny explained between 63-68% of the variation in birds and 70-79% of the variation in mammals (Tables S2, S4). Bird and mammal granulocytes and mammalian lymphocyte cell proportion scaling patterns were consistent in direction (but not magnitude) with previous analyses of cell concentrations (Downs et al., 2020; Ruhs et al., 2020). However, bird lymphocyte proportions were hypometric here whereas no evidence of allometry in lymphocyte concentrations was previously reported (Ruhs et al., 2020).

### Exercise 2: direct comparisons of allometries among taxa

When bat lymphocyte proportions were directly compared to those of birds and non-volant mammals, we found equivalent support for the mass model (model 5; Table 2, Fig. 2), the model with independent effects of mass and taxon (model 6; ΔDIC=0.67), and the model in which allometries differed between taxa (model 7; ΔDIC=3.02). For the mass model (model 5), phylogeny explained a large proportion of the variance (~70%), but the addition of mass increased explanatory power by 16% (marginal *R*^2^). For model 6, the addition of mass and taxon increased explanatory power by an additional 13% (marginal *R*^2^). When examining models that compared the lymphocytes among taxa (models 3, 4, 6, 7), bats were different from non-volant mammals but not birds (Table S6). Because the slope for bat lymphocyte proportions and body mass was indistinguishable from zero (from model 5, *b,* CI = −0.08, −0.09:-0.07; exercise 1), this effect was largely driven by differences in the intercept between taxa (−0.3 in bats; −0.14 in birds; −0.09 in mammals). In other words, mean lymphocyte proportions across all body sizes were lower in bats than other mammals.

**Table 2.** Best-fit models predicting circulating leukocyte concentrations in all species (bats, birds and non-volant mammals). Models test for the effects of body mass and taxon on log_10_-transformed lymphocyte and granulocyte proportions (exercise 2). Top models are bolded.

| Model | DIC | ΔDIC | $\lambda$ (unadjusted)<br>[95% CI] | Marginal $R^2$<br>[95% CI] | Conditional $R^2$<br>[95% CI] |
| --- | --- | --- | --- | --- | --- |
| <u>Lymphocytes</u> |  |  |  |  |  |
| 3. $\beta_0$ | -1163.31 | 96.62 | 0.86<br>[0.78:0.92] | | 0.86<br>[0.78:0.92] |
| 4. $\beta_0 + \beta_1 \times \text{taxon}$ | -1164.48 | 95.45 | 0.83<br>[0.42:0.92] | 0.005<br>[0.0001:0.5] | 0.89<br>[0.81:0.95] |
| **5. $\beta_0 + \beta_1 \times \log_{10}(\text{mass})$** | **-1259.93** | **0** | **0.7**<br>**[0.56:0.79]** | **0.16**<br>**[0.09:0.22]** | **0.84**<br>**[0.77:0.9]** |
| **6. $\beta_0 + \beta_1 \times \log_{10}(\text{mass})$**<br>**+ $\beta_2 \times \text{taxon}$** | **-1259.26** | **0.67** | **0.68**<br>**[0.34:0.8]** | **0.13**<br>**[0.06:0.54]** | **0.85**<br>**[0.75:0.93]** |
| **7. $\beta_0 + \beta_1 \times \log_{10}(\text{mass})$**<br>**+ $\beta_2 \times \text{taxon} + \beta_3 \times$**<br>**$\log_{10}(\text{mass}) \times \text{taxon}$** | **-1256.91** | **3.02** | **0.69**<br>**[0.37:0.81]** | **0.13**<br>**[0.06:0.52]** | **0.85**<br>**[0.75:0.93]** |
| <u>Granulocytes</u> |  |  |  |  |  |
| 3. $\beta_0$ | -1395.22 | 78.31 | 0.88<br>[0.8-0.92] | | 0.88<br>[0.8:0.92] |
| 4. $\beta_0 + \beta_1 \times \text{taxon}$ | -1395.46 | 78.07 | 0.81<br>[0.44:0.9] | 0.03<br>[1.2e-4:0.49] | 0.9<br>[0.82:0.96] |
| 5. $\beta_0 + \beta_1 \times \log_{10}(\text{mass})$ | -1465.85 | 7.68 | 0.71<br>[0.58:0.82] | 0.12<br>[0.06:0.19] | 0.83<br>[0.76:0.9] |
| 6. $\beta_0 + \beta_1 \times \log_{10}(\text{mass})$<br>+ $\beta_2 \times \text{taxon}$ | -1466.3 | 7.23 | 0.71<br>[0.37:0.82] | 0.1<br>[0.04:0.54] | 0.85<br>[0.77:0.94] |
| **7. $\beta_0 + \beta_1 \times \log_{10}(\text{mass})$**<br>**+ $\beta_2 \times \text{taxon} + \beta_3 \times$**<br>**$\log_{10}(\text{Mass}) \times \text{taxon}$** | **-1473.53** | **0** | **0.7**<br>**[0.38:0.83]** | **0.11**<br>**[0.05:0.52]** | **0.87**<br>**[0.77:0.94]** |

Granulocyte proportions, however, were best explained by the model in which intercepts of allometries varied among taxa (model 7); however, this was driven by a universal hypermetric scaling slope across all species (*b*, CI= 0.06, 0.05:07). When examining models comparing granulocyte proportions among taxa (models 3, 4, 6, 7), bat slopes and intercepts were not different from non-volant mammals or birds (Table S7). Phylogeny explained 70% of the total variation. Inclusion of mass and taxa increased explanatory power by 11% (marginal *R^2^*). Therefore, phylogeny explains the majority of the variance in the proportions of both cell types, but body mass informs a moderate percentage of this variation, especially compared to taxon alone (marginal *R^2^*=0.03).

## Discussion

Although bats and birds represent two independent evolutionary origins of flight (Rayner, 1988), both are flying endotherms. We therefore predicted they would be subject to similar selective pressures on their physiology that would ultimately impact their immune system (McGuire and Guglielmo, 2009). Here, we quantified the scaling relationships for proportions of two leukocyte types across 60 species of bats and compared these patterns across other vertebrate endotherms. Broad, comparative analyses of immunity between bats and other taxa are rare (Becker et al., 2019; Shaw et al., 2017), and allometric studies provide a powerful framework for systematically comparing such data across species (Downs et al., 2020, Ruhs et al., 2020). Therefore, our analyses aimed to shed light on bat cellular immunity, generally, and the role of flight in shaping immune scaling relationships. When we examined body mass effects on bat cell proportions (posterior means from exercise 1; Fig. 2), we found little evidence for allometric scaling of either cell type across bat species. When comparing across taxa (exercise 2), however, bat lymphocyte proportions (represented by the intercept) more closely resembled those of birds than non-volant mammals, as the confidence intervals for bats overlapped with those for birds, but not for non-volant mammals. However, bat neutrophil proportions were not distinguishable from birds or non-volant mammals, and all taxa tended to scale hypermetrically. Therefore, our results support the idea that physiological alterations to facilitate flight may not explain the allometry of cell proportions in bats and birds. Our inability to comparatively distinguish bats from other endotherms in exercise 2 is likely driven by the large amount of variation in the bat data, which are from wild populations, and thus complicates identifying allometric patterns; however, as the diversity of leukocyte data from captive bats are low, this comparative analysis can still shed light on flight in shaping immune scaling patterns and help generate predictions for future comparisons. Below we focus on the results from exercise 1 to discuss scaling patterns for each taxon and the (likely) lack of allometry in bats. We then address immunological similarities between bats, birds, and non-volant mammals and what this might mean for pathogen tolerance to motivate future comparative research.

### Are bats immunologically different?

We predicted that bats, like birds, might need a disproportionately large amount of broadly protective cells as they are more likely to be exposed to more and diverse parasites than non-volant species, especially at larger body sizes. However, we found little evidence for intercept or slope differences between bats and non-volant species or allometric scaling in cell proportions of bats. Interestingly, while bats did not show any allometries when analyzed alone (exercise 1), in exercise 2, where cell proportions were compared across taxa, bat lymphocyte proportions more closely resembled birds than non-volant mammals (taxa models panel; Fig. 2). The (likely) lack of scaling, or possible isometry, in bats is intriguing given the evident allometric scaling patterns observed previously for birds and across primarily terrestrial mammals, both for cell concentrations and the proportions analyzed here (Ruhs et al., 2020).

Importantly, contrasting leukocyte patterns between bats and other taxa could be influenced by the bat data being from wild populations and the bird/mammal data being from captive populations. Wild populations are inherently more immunologically variable (Viney & Riley, 2014), which is reflected in the large confidence intervals around our posterior mean estimates in bats. Many factors can influence wild-derived variation, but wild and captive animals can especially differ in pathogen exposure and stressors that can affect leukocyte composition (Davis et al., 2008; Herrera et al., 2019). Alternatively, the absence of allometry in our bat data (when analyzed alone in exercise 1), which are from wild populations, might be more likely to reflect true developmental and environmental pressures on these species. In fact, another study demonstrated that the variation in cell proportions from wild species did not influence scaling patterns, as wild and captive birds both displayed similar allometries (Ruhs et al., *unpublished data).* Similarly, comparison of wild and captive rodents also revealed no differences between scaling patterns of total leukocytes or neutrophil counts, despite captive animals having higher mean lymphocyte counts than wild animals (Tian et al., 2015). Taken in sum, the lack of scaling patterns found here are unlikely driven by variation in wild bat cell proportions.

The small differences in bat versus other taxa cell proportion allometries support similar efforts to understand constitutive expression of other aspects of the bat immune system. For example, comparative genomic analyses show several unusual immunological aspects of bats, such as high constitutive expression of type I interferons and dampened inflammation (Ahn et al., 2019; Pavlovich et al., 2018; Zhou et al., 2016). Although our results are limited to proportions of two key white blood cells, the inability to distinguish bats from birds (for lymphocytes) and bats from all other taxa (for neutrophils) suggest there may be other ecological explanations (e.g. not flight-related) for the cell proportion scaling patterns.

Given the results from exercise 1, it is instead possible that the bat data are representative of isometry, such that species require the same proportions of leukocytes across body size. Isometry could be driven by certain distinct aspects of bat biology. A plausible explanation could involve their slow life-history strategy. Bats, even more so than birds, are long-lived such that selection for safety and disease risk-reduction is likely prioritized to accommodate longevity, across all body masses. Although body mass affects longevity in bats, other factors such as hibernation, cave use, and latitude all have effects of similar magnitude on lifespan as mass (Wilkinson and Adams 2019), which may complicate detecting mass effects on physiology or morphology linked to lifespan. For example, high sociality and gregariousness of many bat species facilitates contact during roosting that could increase pathogen exposure (Kerth, 2008; Kunz, 1982; Webber et al., 2017) and would be equal across body size. Although it is most likely that the bat data here represent a lack of scaling, these alternative explanations support the possibility that our bat data might represent isometry of lymphocytes when directly comparing bats to other taxa.

### Future directions

Increased spillover of zoonotic viruses, such as henipaviruses and coronaviruses, has renewed public and scientific interest in whether bats are immunologically unique reservoir hosts (Brook and Dobson, 2015; Halpin et al., 2011; Li et al., 2005; Luis et al., 2013). Investigating allometric scaling patterns of immunological features, such as leukocytes, could shed light on the physiological traits that impact host ability to tolerate virulent pathogens. We here demonstrate some differences in the scaling patterns of innate immune cell proportions between taxa of endotherms; however, we did not observe substantial effects of body size on cell proportions in bats. It is important to note that it is likely more difficult to find allometric patterns of cell proportions using the methods employed here compared to the previous discovery of hypermetric scaling of cell concentrations (Downs et al., 2020; Ruhs et al., 2020); therefore, future studies should aim to measure cell concentrations, which are much more insightful for immunological patterns. Our sample also represents only a small fraction of bat diversity (about 4% of the >1400 species; Simmons and Cirranello, 2020), although our data do span the body mass continuum of extant bat species. To enhance our ability to examine relevant patterns, we encourage greater breadth of immunological studies across the bat phylogeny, specifically within the bat clades characterized by relatively larger body sizes (e.g., Pteropodidae).

Lastly, we focused on cell proportion allometry and the potential for body mass alone to explain immunological differences among species (Downs et al., 2019). Flying endotherms can vary in other ecological traits besides body mass that also shape pathogen exposure and immune investment, such as diet, coloniality, and roost type (Minias, Whittingham, & Dunn, 2017; Schneeberger et al., 2013). To address such trait comparisons across equal ecological context (i.e., avoiding captive-wild contrasts), future comparative studies of wild bats, birds, and non-volant mammals of similar body masses could help to confirm the patterns observed here and further differentiate evolutionary effects from those of flight and other aspects of life history on immune defense.

## Acknowledgments

For assistance with field logistics and permits, we thank Mark Howells, Neil Duncan, and the staff of the Lamanai Field Research Center. We also thank the many colleagues who helped net bats during 2017 and 2018 bat research in Belize as well as Ellen Chinchilli and Grace Carey for help compiling bird and mammal data. Lastly, we thank members of the Martin lab at the University of South Florida, the Ketterson lab at Indiana University, and anonymous reviewers for constructive feedback.

## Competing interests

The authors declare no competing interests.

## Author contributions

DJB collected field data; ECR and SO collected literature data; SO, HFD, and OO analyzed blood smears; ECR performed the statistical analyses; and MBF, NBS, LBM, and CJD provided logistical and funding support. ECR and DJB wrote the manuscript with input from coauthors, and all authors gave final approval for publication.

## Funding

This work was funded by the ARCS Foundation and Explorer’s Club (DJB), American Museum of Natural History (Theodore Roosevelt Memorial Fund to DJB, Taxonomic Mammalogy Fund to NBS), and National Science Foundation (IOS 1656618 to LBM, IOS 1656551 to CJD).

## Data accessibility

Bat leukocyte data and endotherm metadata will be deposited in Dryad Digital Repository. Captive bird and mammal white blood cell data are available from Species360.

## References

Ahn, M., Anderson, D. E., Zhang, Q., Tan, C. W., Lim, B. L., Luko, K., … Wang, L.-F. (2019). Dampened NLRP3-mediated inflammation in bats and implications for a special viral reservoir host. Nature Microbiology, 4(5), 789–799.

Albery, G. F., Watt, K. A., Keith, R., Morris, S., Morris, A., Kenyon, F., … Pemberton, J. M. (n.d.). Reproduction has different costs for immunity and parasitism in a wild mammal. doi: 10.1101/472597

Altizer, S., Bartel, R., & Han, B. A. (2011). Animal migration and infectious disease risk. Science, 331(6015), 296–302.

Amman, B. R., Jones, M. E. B., Sealy, T. K., Uebelhoer, L. S., Schuh, A. J., Bird, B. H., … Towner, J. S. (2015). Oral shedding of Marburg virus in experimentally infected Egyptian fruit bats (Rousettus aegyptiacus). Journal of Wildlife Diseases, 51(1), 113–124.

Baker, M. L., Schountz, T., & Wang, L.-F. (2013). Antiviral immune responses of bats: a review. Zoonoses and Public Health, 60(1), 104–116.

Banyard, A. C., Hayman, D., Johnson, N., McElhinney, L., & Fooks, A. R. (2011). Bats and lyssaviruses. Advances in Virus Research, 79, 239–289.

Becker, D. J., Czirják, G. Á., Rynda-Apple, A., & Plowright, R. K. (2019). Handling Stress and Sample Storage Are Associated with Weaker Complement-Mediated Bactericidal Ability in Birds but Not Bats. Physiological and Biochemical Zoology, Vol. 92, pp. 37–48. doi: 10.1086/701069

Becker, D. J., Czirják, G. Á., & Volokhov, D. V. (2018). Livestock abundance predicts vampire bat demography, immune profiles and bacterial infection risk. Of the Royal…. Retrieved from https://royalsocietypublishing.org/doi/abs/10.1098/rstb.2017.0089

Brace, A. J., Lajeunesse, M. J., Ardia, D. R., Hawley, D. M., Adelman, J. S., Buchanan, K. L., … Martin, L. B. (2017). Costs of immune responses are related to host body size and lifespan. Journal of Experimental Zoology. Part A, Ecological and Integrative Physiology, 327(5), 254–261.

Brigham, R. M. (1991). Prey Detection, Dietary Niche Breadth, and Body Size in Bats: Why are Aerial Insectivorous Bats so Small? The American Naturalist, 137(5), 693–703.

Brook, C. E., Boots, M., Chandran, K., Dobson, A. P., Drosten, C., Graham, A. L., … van Leeuwen, A. (2020). Accelerated viral dynamics in bat cell lines, with implications for zoonotic emergence. eLife, 9. doi: 10.7554/eLife.48401

Brook, C. E., & Dobson, A. P. (2015). Bats as “special”reservoirs for emerging zoonotic pathogens. Trends in Microbiology, 23(3), 172–180.

Brown, J. H., Gillooly, J. F., Allen, A. P., & Savage, V. M. (2004). Toward a metabolic theory of ecology. Ecology.

Calder, W. A. (1996). Size, Function, and Life History. Courier Corporation.

Carter, G., & Leffer, L. (2015). Social Grooming in Bats: Are Vampire Bats Exceptional? PloS One, 10(10), e0138430.

Clayton, D. H., Koop, J. A. H., Harbison, C. W., Moyer, B. R., & Bush, S. E. (2010). How Birds Combat Ectoparasites. The Open Ornithology Journal, Vol. 3, pp. 41–71. doi: 10.2174/1874453201003010041

Cohn, M., & Langman, R. E. (1990). The Protection: The Unit of Humoral Immunity Selected by Evolution. Immunological Reviews, 115(1), 7–147.

Crichton, E. G., & Krutzsch, P. H. (2000). Reproductive Biology of Bats. Academic Press.

Davis, A. K., Maney, D. L., & Maerz, J. C. (2008). The use of leukocyte profiles to measure stress in vertebrates: a review for ecologists. Functional Ecology, 22(5), 760–772.

Delius, J. D. (1988). Preening and associated comfort behavior in birds. Annals of the New York Academy of Sciences, 525, 40–55.

Dingli, D., & Pacheco, J. M. (2006). Allometric scaling of the active hematopoietic stem cell pool across mammals. PloS One, 1, e2.

Downs, C. J., Dochtermann, N. A., Ball, R., Klasing, K. C., & Martin, L. B. (2020). The Effects of Body Mass on Immune Cell Concentrations of Mammals. The American Naturalist, 195(1), 107–114.

Downs, C. J., Schoenle, L. A., Han, B. A., Harrison, J. F., & Martin, L. B. (2019). Scaling of Host Competence. Trends in Parasitology, 35(3), 182–192.

Dumont, E. R. (2010). Bone density and the lightweight skeletons of birds. Proceedings. Biological Sciences / The Royal Society, 277(1691), 2193–2198.

Dunning, J. B., Jr. (2007). CRC Handbook of Avian Body Masses. CRC Press.

Feduccia, A., & Feduccia, A. (1999). The Origin and Evolution of Birds. Yale University Press.

Frick, W. F., Puechmaille, S. J., Hoyt, J. R., Nickel, B. A., Langwig, K. E., Foster, J. T., … Others. (2015). Disease alters macroecological patterns of North American bats. Global Ecology and Biogeography: A Journal of Macroecology, 24(7), 741–749.

Gunnell, G. F., & Simmons, N. B. (2012). Evolutionary History of Bats: Fossils, Molecules and Morphology. Cambridge University Press.

Guth, S., Visher, E., Boots, M., & Brook, C. E. (2019). Host phylogenetic distance drives trends in virus virulence and transmissibility across the animal–human interface. Philosophical Transactions of the Royal Society of London. Series B, Biological Sciences, 374(1782), 20190296.

Hadfield, J. D., & Others. (2010). MCMC methods for multi-response generalized linear mixed models: the MCMCglmm R package. Journal of Statistical Software, 33(2), 1–22.

Halpin, K., Hyatt, A. D., Fogarty, R., Middleton, D., Bingham, J., Epstein, J. H., … Henipavirus Ecology Research Group. (2011). Pteropid bats are confirmed as the reservoir hosts of henipaviruses: a comprehensive experimental study of virus transmission. The American Journal of Tropical Medicine and Hygiene, 85(5), 946–951.

Hamm, P. S., Caimi, N. A., Northup, D. E., Valdez, E. W., Buecher, D. C., Dunlap, C. A., … Porras-Alfaro, A. (2017). Western Bats as a Reservoir of Novel Streptomyces Species with Antifungal Activity. Applied and Environmental Microbiology, 83(5). doi: 10.1128/AEM.03057-16

Han, B. A., Park, A. W., Jolles, A. E., & Altizer, S. (2015). Infectious disease transmission and behavioural allometry in wild mammals. Journal of Animal Ecology, Vol. 84, pp. 637–646. doi: 10.1111/1365-2656.12336

Harrison, J. F. (2017). Do Performance–Safety Tradeoffs Cause Hypometric Metabolic Scaling in Animals? Trends in Ecology & Evolution, 32(9), 653–664.

Hatten, B. A., Lutskus, J. H., & Sulkin, S. E. (1973). A serologic comparison of bat complements. The Journal of Experimental Zoology, 186(2), 193–206.

Healy, K., Guillerme, T., Finlay, S., Kane, A., Kelly, S. B. A., McClean, D., … Cooper, N. (2014). Ecology and mode-of-life explain lifespan variation in birds and mammals. Proceedings. Biological Sciences / The Royal Society, 281(1784), 20140298.

Herrera, J. P., Chakraborty, D., Rushmore, J., Altizer, S., & Nunn, C. (2019). The changing ecology of primate parasites: Insights from wild captive comparisons. American Journal of Primatology, 81(7), 29.

Herrera, J. P., Duncan, N., Clare, E., Brock Fenton, M., & Simmons, N. (2018). Disassembly of Fragmented Bat Communities in Orange Walk District, Belize. Acta Chiropterologica, Vol. 20, p. 147. doi: 10.3161/15081109acc2018.20.1.011

Housworth, E. A., Martins, E. P., & Lynch, M. (2004). The phylogenetic mixed model. The American Naturalist, 163(1), 84–96.

Ingala, M. R., Simmons, N. B., & Perkins, S. L. (2018). Bats Are an Untapped System for Understanding Microbiome Evolution in Mammals. mSphere, 3(5). doi: 10.1128/mSphere.00397-18

Jimenez, A. G., O’Connor, E. S., Tobin, K. J., Anderson, K. N., Winward, J. D., Fleming, A., … Downs, C. J. (2019). Does Cellular Metabolism from Primary Fibroblasts and Oxidative Stress in Blood Differ between Mammals and Birds? The (Lack-thereof) Scaling of Oxidative Stress. Integrative and Comparative Biology, 59(4), 953–969.

Jones, G., & Teeling, E. C. (2006). The evolution of echolocation in bats. Trends in Ecology & Evolution, 21(3), 149–156.

Jones, T., Myers, P., & Espinosa, R. (1997). Animal Diversity Web. Media.

Kacprzyk, J., Hughes, G. M., Palsson-McDermott, E. M., Quinn, S. R., Puechmaille, S. J., O’neill, L. A. J., & Teeling, E. C. (2017). A potent anti-inflammatory response in bat macrophages may be linked to extended longevity and viral tolerance. Acta Chiropterologica / Museum and Institute of Zoology, Polish Academy of Sciences, 19(2), 219–228.

Kerth, G. (2008). Causes and Consequences of Sociality in Bats. Bioscience, 58(8), 737–746.

Kessler, M. K., Becker, D. J., & Peel, A. J. (2018). Changing resource landscapes and spillover of henipaviruses. Annals of the New York Academy of Sciences. Retrieved from https://www.ncbi.nlm.nih.gov/pmc/articles/PMC6778453/

Kleiber, M., & Others. (1932). Body size and metabolism. Hilgardia, 6(11), 315–353.

Kunz, T. H. (1982). Roosting Ecology of Bats. In T. H. Kunz (Ed.), Ecology of Bats (pp. 1–55). Boston, MA: Springer US.

Kunz, T. H., Braun de Torrez, E., Bauer, D., Lobova, T., & Fleming, T. H. (2011). Ecosystem services provided by bats. Annals of the New York Academy of Sciences, 1223, 1–38.

Kunz, T. H., & Kurta, A. (1987). Size of bats at birth and maternal investment during pregnancy. Symposia of the Zoological Society of London, 57, 79–106.

Lanier, L. L. (2013). Shades of grey — the blurring view of innate and adaptive immunity. Nature Reviews Immunology, Vol. 13, pp. 73–74. doi: 10.1038/nri3389

Lee, K. A. (2006). Linking immune defenses and life history at the levels of the individual and the species. Integrative and Comparative Biology, 46(6), 1000–1015.

Letunic, I., 2015. Phylot: Phylogenetic Tree Generator. [online] Phylot.biobyte.de. Available at: <http://phylot.biobyte.de/>.

Levesque, DL, JG Boyles, CJ Downs, and AM Breit. In Press. High body temperature is an unlikely cause of high tolerance in bats. Journal of Wildlife Diseases.

Li, W., Shi, Z., Yu, M., Ren, W., Smith, C., Epstein, J. H., … Wang, L.-F. (2005). Bats are natural reservoirs of SARS-like coronaviruses. Science, 310(5748), 676–679.

Luis, A. D., Hayman, D. T. S., O’Shea, T. J., Cryan, P. M., Gilbert, A. T., Pulliam, J. R. C., … Webb, C. T. (2013). A comparison of bats and rodents as reservoirs of zoonotic viruses: are bats special? Proceedings. Biological Sciences / The Royal Society, 280(1756), 20122753.

Mandl, J. N., Schneider, C., Schneider, D. S., & Baker, M. L. (2018). Going to Bat(s) for Studies of Disease Tolerance. Frontiers in Immunology, 9, 2112.

Martin, L. B., Weil, Z. M., & Nelson, R. J. (2008). Seasonal changes in vertebrate immune activity: mediation by physiological trade-offs. Philosophical Transactions of the Royal Society of London. Series B, Biological Sciences, 363(1490), 321–339.

Maurer, B. A., Alroy, J., Brown, J. H., Dayan, T., Enquist, B., Morgan Ernest, S. K., … Willig, M. R. (2004). Similarities in body size distributions of small-bodied flyingvertebrates. Evolutionary Ecology Research, 6, 783.

McGuire, L. P., & Guglielmo, C. G. (2009). What Can Birds Tell Us about the Migration Physiology of Bats? Journal of Mammalogy, 90(6), 1290–1297.

Minias, P., Whittingham, L. A., & Dunn, P. O. (2017). Coloniality and migration are related to selection on MHC genes in birds. Evolution; International Journal of Organic Evolution, 71(2), 432–441.

Mollentze, N., & Streicker, D. G. (2020). Viral zoonotic risk is homogenous among taxonomic orders of mammalian and avian reservoir hosts. Proceedings of the National Academy of Sciences, Vol. 117, pp. 9423–9430. doi: 10.1073/pnas.1919176117

Muijres, F. T., Johansson, L. C., Bowlin, M. S., Winter, Y., & Hedenström, A. (2012). Comparing aerodynamic efficiency in birds and bats suggests better flight performance in birds. PloS One, 7(5). Retrieved from https://www.ncbi.nlm.nih.gov/pmc/articles/pmc3356262/

Munshi-South, J., & Wilkinson, G. S. (2010). Bats and birds: Exceptional longevity despite high metabolic rates. Ageing Research Reviews, 9(1), 12–19.

Nakagawa, S., & Schielzeth, H. (2013). A general and simple method for obtaining R2 from generalized linear mixed-effects models. Methods in Ecology and Evolution /British Ecological Society, 4(2), 133–142.

Ogburn, C. E., Austad, S. N., Holmes, D. J., Kiklevich, J. V., Gollahon, K., Rabinovitch, P. S., & Martin, G. M. (1998). Cultured renal epithelial cells from birds and mice: enhanced resistance of avian cells to oxidative stress and DNA damage. The Journals of Gerontology. Series A, Biological Sciences and Medical Sciences, 53(4), B287–B292.

Ogburn, C. E., Carlberg, K., Ottinger, M. A., Holmes, D. J., Martin, G. M., & Austad, S. N. (2001). Exceptional cellular resistance to oxidative damage in long-lived birds requires active gene expression. The Journals of Gerontology. Series A, Biological Sciences and Medical Sciences, 56(11), B468–B474.

O’Shea, T. J., Cryan, P. M., Cunningham, A. A., Fooks, A. R., Hayman, D. T. S., Luis, A. D., … Wood, J. L. N. (2014). Bat flight and zoonotic viruses. Emerging Infectious Diseases, 20(5), 741–745.

Paradis, E., Claude, J., & Strimmer, K. (2004). APE: Analyses of Phylogenetics and Evolution in R language. Bioinformatics, 20(2), 289–290.

Parr, C. S., Wilson, N., Leary, P., Schulz, K. S., Lans, K., Walley, L., … Corrigan, R. J., Jr. (2014). The Encyclopedia of Life v2: Providing Global Access to Knowledge About Life on Earth. Biodiversity Data Journal, (2), e1079.

Pavlovich, S. S., Lovett, S. P., Koroleva, G., Guito, J. C., Arnold, C. E., Nagle, E. R., … Palacios, G. (2018). The Egyptian Rousette Genome Reveals Unexpected Features of Bat Antiviral Immunity. Cell, 173(5), 1098–1110.e18.

Peel, A. J., Wells, K., Giles, J., Boyd, V., Burroughs, A., Edson, D., … Clark, N. (2019). Synchronous shedding of multiple bat paramyxoviruses coincides with peak periods of Hendra virus spillover. Emerging Microbes & Infections, 8(1), 1314–1323.

Plowright, R. K., Field, H. E., Smith, C., Divljan, A., Palmer, C., Tabor, G., … Foley, J. E. (2008). Reproduction and nutritional stress are risk factors for Hendra virus infection in little red flying foxes (Pteropus scapulatus). Proceedings. Biological Sciences / The Royal Society, 275(1636), 861–869.

Podlutsky, A. J., Khritankov, A. M., Ovodov, N. D., & Austad, S. N. (2005). A new field record for bat longevity. The Journals of Gerontology. Series A, Biological Sciences and Medical Sciences, 60(11), 1366–1368.

Rayner, J. M. V. (1988). The evolution of vertebrate flight. Biological Journal of the Linnean Society. Linnean Society of London, 34(3), 269–287.

Reid, F. (1997). A field guide to the mammals of Central America and Southeast México: Oxford University Press. New York, 334.

Richards, S. A. (2005). Testing ecological theory using the information-theoretic approach: examples and cautionary results. Ecology, 86(10), 2805–2814.

Robert M. R. Barclay. (1994). Constraints on Reproduction by Flying Vertebrates: Energy and Calcium. The American Naturalist, 144(6), 1021–1031.

Ruhs, E. C., Martin, L. B., & Downs, C. J. (2020). The impacts of body mass on immune cell concentrations in birds. doi: 10.1101/2020.04.23.057794

Schmidt-Nielsen, K. (1972). Locomotion: energy cost of swimming, flying, and running. Science, 177(4045), 222–228.

Schmidt-Nielsen, K., & Knut, S.-N. (1984). Scaling: Why is Animal Size So Important? Cambridge University Press.

Schneeberger, K., Czirják, G. Á., & Voigt, C. C. (2013). Measures of the constitutive immune system are linked to diet and roosting habits of neotropical bats. PloS One, 8(1), e54023.

Seltmann, A., Corman, V. M., Rasche, A., Drosten, C., Czirják, G. Á., Bernard, H., … Voigt, C. C. (2017). Seasonal Fluctuations of Astrovirus, But Not Coronavirus Shedding in Bats Inhabiting Human-Modified Tropical Forests. EcoHealth, 14(2), 272–284.

Seltmann, A., Czirják, G. Á., Courtiol, A., Bernard, H., Struebig, M. J., & Voigt, C. C. (2017). Habitat disturbance results in chronic stress and impaired health status in forest-dwelling paleotropical bats. Conservation Physiology, 5(1), cox020.

Shaw, A. E., Hughes, J., Gu, Q., Behdenna, A., Singer, J. B., Dennis, T., … Palmarini, M. (2017). Fundamental properties of the mammalian innate immune system revealed by multispecies comparison of type I interferon responses. PLoS Biology, 15(12), e2004086.

Shawkey, M. D., Pillai, S. R., & Hill, G. E. (2003). Chemical warfare? Effects of uropygial oil on feather-degrading bacteria. Journal of Avian Biology, Vol. 34, pp. 345–349. doi: 10.1111/j.0908-8857.2003.03193.x

Sikes, R. S., & Gannon, W. L. (2011). Guidelines of the American Society of Mammalogists for the use of wild mammals in research. Journal of Mammalogy, 92(1), 235–253.

Simmons, N. B., & Cirranello, A. L. (2020). Bat Species of the World: A taxonomic and geographic database. Accessed on, 7(10), 2020.

Speakman, J. R. (2008). The physiological costs of reproduction in small mammals. Philosophical Transactions of the Royal Society of London. Series B, Biological Sciences, 363(1490), 375–398.

Stanley, S. M. (1973). AN EXPLANATION FOR COPE’S RULE. Evolution; International Journal of Organic Evolution, 27(1), 1–26.

Stockmaier, S., Dechmann, D. K. N., Page, R. A., & O’Mara, M. T. (2015). No fever and leucocytosis in response to a lipopolysaccharide challenge in an insectivorous bat. Biology Letters, 11(9), 20150576.

Swartz, S., Iriarte-Diaz, J., Riskin, D., Tian, X., Song, A., & Breuer, K. (2007). Wing Structure and the Aerodynamic Basis of Flight in Bats. In Aerospace Sciences Meetings. 45th AIAA Aerospace Sciences Meeting and Exhibit. American Institute of Aeronautics and Astronautics.

Tacutu, R., Craig, T., Budovsky, A., Wuttke, D., Lehmann, G., Taranukha, D., … de Magalhães, J. P. (2013). Human Ageing Genomic Resources: integrated databases and tools for the biology and genetics of ageing. Nucleic Acids Research, 41(Database issue), D1027–D1033.

Teeling, E. C., Vernes, S. C., Dávalos, L. M., Ray, D. A., Gilbert, M. T. P., Myers, E., & Bat1K Consortium. (2018). Bat Biology, Genomes, and the Bat1K Project: To Generate Chromosome-Level Genomes for All Living Bat Species. Annual Review of Animal Biosciences, 6, 23–46.

Thomas, S. P. (1975). Metabolism during flight in two species of bats, Phyllostomus hastatus and Pteropus gouldii. The Journal of Experimental Biology, 63(1), 273–293.

Tian, J., Courtiol, A., Schneeberger, K., Greenwood, A. D., & Czirják, G. Á. (2015). Circulating white blood cell counts in captive and wild rodents are influenced by body mass rather than testes mass, a correlate of mating promiscuity. Functional Ecology, 29(6), 823–829.

Tian, X., Iriarte-Diaz, J., Middleton, K., Galvao, R., Israeli, E., Roemer, A., … Breuer, K. (2006). Direct measurements of the kinematics and dynamics of bat flight. Bioinspiration & Biomimetics, 1(4), S10–S18.

Viney, M. E., & Riley, E. M. (2014). From Immunology to Eco-Immunology: More than a New Name. In D. Malagoli & E. Ottaviani (Eds.), Eco-immunology: Evolutive Aspects and Future Perspectives (pp. 1–19). Dordrecht: Springer Netherlands.

Webber, Q. M. R., Fletcher, Q. E., & Willis, C. K. R. (2017). Viral Richness is Positively Related to Group Size, but Not Mating System, in Bats. EcoHealth, 14(4), 652–661.

West, G. B., Brown, J. H., & Enquist, B. J. (2000). Scaling in biology: patterns and processes, causes and consequences. Scaling in Biology, 87–112.

Wiegel, F. W., & Perelson, A. S. (2004). Some scaling principles for the immune system. Immunology and Cell Biology, 82(2), 127–131.

Wilkinson, G. S., & Adams, D. M. (2019). Recurrent evolution of extreme longevity in bats. Biology Letters, 15(4), 20180860.

Wilkinson, G. S., & South, J. M. (2002). Life history, ecology and longevity in bats. Aging Cell, 1(2), 124–131.

Williamson, M. M., Hooper, P. T., Selleck, P. W., Westbury, H. A., & Slocombe, R. F. (2000). Experimental hendra virus infectionin pregnant guinea-pigs and fruit Bats (Pteropus poliocephalus). Journal of Comparative Pathology, 122(2-3), 201–207.

Wilman, H., Belmaker, J., Simpson, J., De La Rosa, C., Rivadeneira, M. M., & Jetz, W. (2014). EltonTraits 1.0: Species-level foraging attributes of the world’s birds and mammals: Ecological Archives E095-178. Ecology, 95(7), 2027–2027.

Xie, J., Li, Y., Shen, X., Goh, G., Zhu, Y., Cui, J., … Zhou, P. (2018). Dampened STING-Dependent Interferon Activation in Bats. Cell Host & Microbe, 23(3), 297–301.e4.

Zhang, G., Cowled, C., Shi, Z., Huang, Z., Bishop-Lilly, K. A., Fang, X., … Wang, J. (2013). Comparative analysis of bat genomes provides insight into the evolution of flight and immunity. Science, 339(6118), 456–460.

Zhou, P., Tachedjian, M., Wynne, J. W., Boyd, V., Cui, J., Smith, I., … Baker, M. L. (2016). Contraction of the type I IFN locus and unusual constitutive expression of IFN-α in bats. Proceedings of the National Academy of Sciences of the United States of America, 113(10), 2696–2701.

ZIMS: Zoo aquarium animal management software - Species360. Retrieved April, 2019, from Species360 website: https://www.species360.org/products-services/zoo-aquarium-animal-management-software-2/

